# Effect of flavonoids hydroxygenkwanin on vascular smooth muscle cell proliferation, migration, and neointimal formation

**DOI:** 10.1101/2022.12.20.521220

**Authors:** Chin-Chuan Chen, Mao-Shin Lin, Pin-Yu Chen, Yann-Lii Leu, Shu-Huei Wang

## Abstract

**Background:** Restenosis and atherosclerosis are chronic inflammatory disease. Abnormal vascular smooth muscle cell (VSMC) proliferation and migration play crucial roles in neointimal hyperplasia and restenosis progression in response to stimulation with various inflammatory cytokines, such as platelet-derived growth factor-BB (PDGF-BB) and tumour necrosis factor-α (TNF-α). Hydroxygenkwanin (HGK) exerts remarkable anti-inflammatory, antitumour, antiproliferative and antimigratory effects. The aim of the study was to evaluate and elucidate the therapeutic effect and regulatory mechanism of HGK on neointimal hyperplasia.

**Methods:** To determine the therapeutic effects of HGK in PDGF-BB- or TNF-α-treated VSMCs, MTT assays, Western blotting analysis, cell cycle analysis, BrdU incorporation assay, wound healing assay and adhesion assay were performed in vitro. A docking assay was also used to elucidate the mechanism underlying the regulatory effect of HGK. Histological and immunohistochemical staining of denuded femoral arteries was conducted to elucidate the therapeutic effect of HGK in an in vivo assay.

**Results:** HGK inhibited the abnormal proliferation, migration, and inflammation of PDGF-BB- or TNF-α-treated VSMCs through regulation of the PDK1/AKT/mTOR pathway. In addition, HGK promoted circulating endothelial progenitor cell (EPC) chemotaxis. In an in vivo assay, HGK dramatically enhanced re-endothelization and reduced neointimal hyperplasia after femoral artery denudation with a guide wire in mice.

**Conclusions:** In the present study, HGK decreased the PDGF-BB- or TNF-α-induced abnormal proliferation, migration and inflammation in VSMCs and improved re-endothelialization and neointimal hyperplasia in denuded femoral arteries. These results provide a novel potential treatment for restenosis in the future.

**Graphic abstract:** HGK decreases VSMC abnormal proliferation, migration and inflammation through PDK1/AKT/mTOR/S6K inhibition and promotes EPC chemotaxis and reendothelialization. HGK is a potential therapeutic candidate for intimal hyperplasia and restenosis.

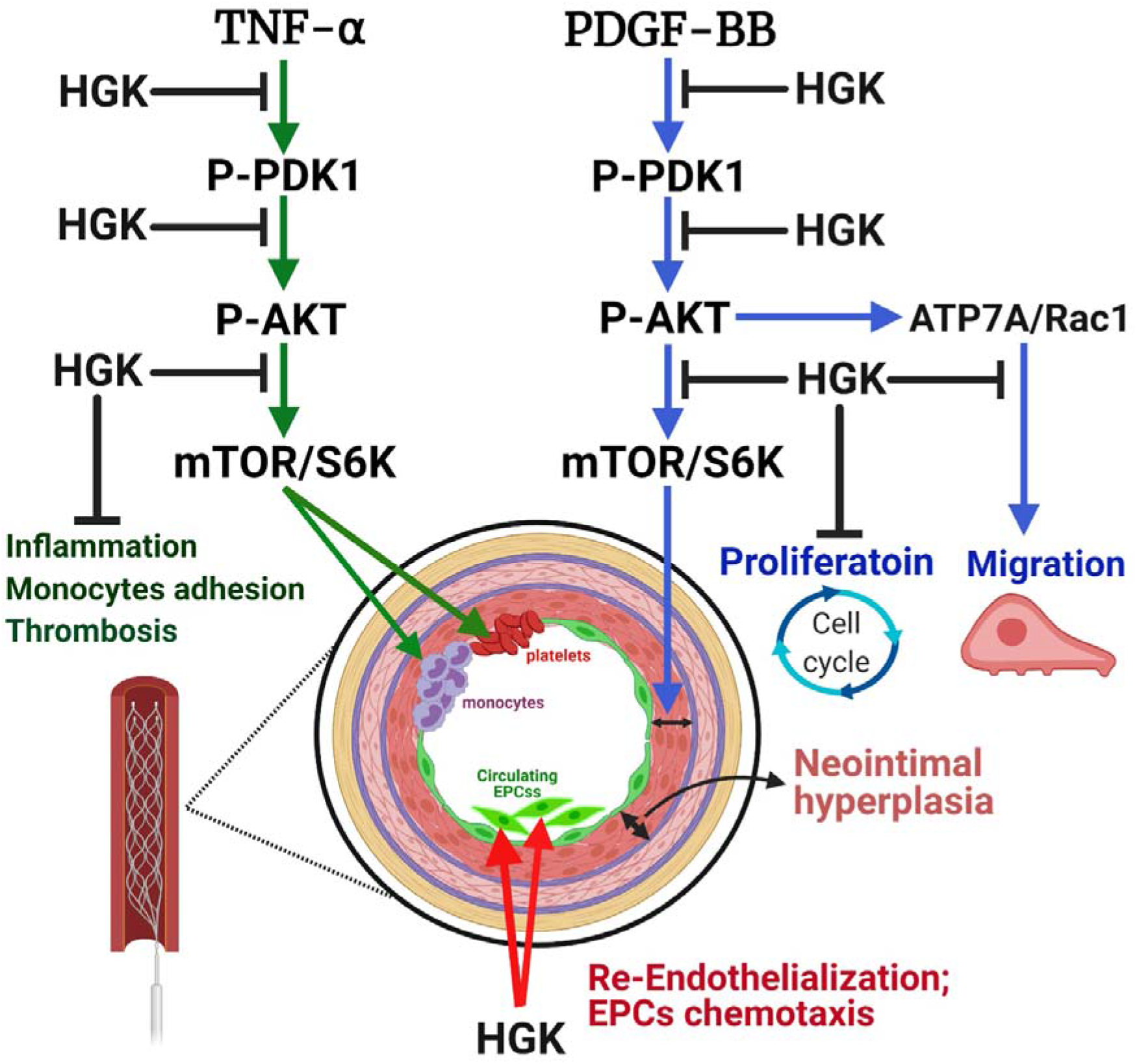

## 1. Background

Restenosis, the renarrowing of a blood vessel after angioplasty, occurs mainly through a process of neointimal hyperplasia and is caused by various events, such as endothelial denudation (ED) or injury, platelet activation, inflammation and abnormal proliferation and migration of vascular smooth muscle cells (VSMCs) (1). Platelet aggregation and activation are induced after endothelial injury and dysfunction and then increase inflammatory cell adhesion, infiltration and interaction with VSMCs (2). The stimulated VSMCs in the tunica media undergo transformation from a contractile phenotype to a synthetic phenotype, and these synthetic VSMCs then substantially exacerbate restenosis progression by promoting proliferation and migration (3). Agents that exert antiproliferative, anti-migratory and anti-inflammatory effects on VSMCs and accelerate reendothelization of the injured endothelium may be developed as new therapeutic agents for neointimal hyperplasia and restenosis.

Hydroxygenkwanin (HGK), a flavonoid, is purified from Daphne genkwa plants. HGK exerts multiple biological effects, including antimicrobial (4), antioxidant (5), and free radical scavenging effects (6), and promotes pigmented hair regeneration (7). In addition, HGK exerts antitumor effects on lung cancer (8), glioma (9), oral squamous cell carcinoma (10), and liver cancer (11) by regulating growth arrest and inhibiting migration. In cardiovascular disease, research has shown that HGK has anti-inflammatory activity and acts as a tissue factor (TF) inhibitor to prevent thrombosis (12). However, the role of HGK in neointimal hyperplasia has not been reported. Therefore, VSMCs and a mouse femoral artery wire-induced injury model were used to test the efficacy of HGK against neointimal hyperplasia and to investigate its regulatory mechanisms in detail. In the present study, HGK exerted antiproliferative, anti-migratory and anti-inflammatory effects, accelerated reendothelization and reduced neointimal hyperplasia.

## 2. Methods

### 2.1 Materials

Rabbit polyclonal IgG antibodies against GAPDH, β-actin, α-tubulin, phospho/total PDK1, phospho-AKT (S473), phospho-AKT (T308), total AKT, total JNK, total ERK, total P38, total mTOR, total S6K, α-SMA, BrdU, PAI-1, and P27, as well as horseradish peroxidase (HRP)-conjugated goat anti-rabbit IgG antibodies and rabbit and mouse IgG isotype control antibodies, were purchased from GeneTex (Irvine, CA, USA). Rabbit polyclonal IgG antibodies against phospho-mTOR, phospho-S6K, phospho-JNK, phospho-ERK, phospho-P38, Cyclin E, and Cyclin D1 were purchased from Cell Signaling Technology (Beverly, MA, USA). Mouse monoclonal IgG antibodies against VCAM-1, ICAM-1, E-selectin, proliferating cell nuclear antigen (PCNA), CDK2 and CDK4 were purchased from Santa Cruz Biotechnology (Dallas, TX, USA). A rabbit polyclonal IgG antibody against ATP7A was purchased from LSBio (Seattle, WA, USA). A rabbit polyclonal IgG antibody against TF was purchased from Bioss (Woburn, MA, USA). Rabbit monoclonal IgG antibodies against MMP-9 and Cx43 and CD31 and a mouse IgG antibody against Rac-1 were purchased from Abcam (Cambridge, UK). Rabbit polyclonal IgG antibodies against Na+/K+-ATPase and CD68 were purchased from Arigo (Hsinchu, Taiwan). MK2206 (an AKT inhibitor), BX795 (a PDK1 inhibitor) and PS48 (a PDK1 activator) were purchased from Selleck Chemicals (Houston, TX, USA). PDGF-BB and TNF-α were purchased from PeproTech (Rocky Hill, NJ, USA). 4’,6-Diamidino-2-phenylindole (DAPI), 3-(4,5-dimethylthiazol-2-yl)-2,5-diphenyl tetrazolium bromide (MTT), phalloidin, elastin and Evans blue dye were purchased from Sigma– Aldrich (St. Louis, MO, USA). 2’,7’-Bis-(2-carboxyethyl)-5-(and-6)-carboxyfluorescein acetoxymethyl ester (BCECF-AM) was purchased from Molecular Probes (Invitrogen, Carlsbad, CA, USA). Fluorescein isothiocyanate (FITC)- and tetramethylrhodamine (TRITC)-conjugated goat anti-mouse and rabbit IgG antibodies were purchased from Jackson ImmunoResearch (West Grove, PA, USA). The ImmPRESS-Alkaline phosphatase polymer kit was purchased from Vector Laboratories (Burlingame, CA, USA). All antibodies and reagents are listed in Supplementary Tables 1 & 2.

### 2.2 Extraction and purification of HGK

HGK was purified from D. genkwa plants. Briefly, the dry flower buds of D. genkwa (5.0 kg) were extracted with MeOH (30 L × 4) and concentrated to produce brown syrup (745.75 g). The syrup was suspended in H2O and successively partitioned with CHCl3 and n-BuOH. The CHCl3 extract (511.19 g) was subjected to column chromatography over silica gel and eluted with CHCl3 and MeOH stepwise gradients to afford fourteen fractions. The seventh fraction was also subjected to column chromatography over silica gel and eluted with n-hexane and acetone step gradients to afford HGK (206.4 mg). HGK (> 95.3% purity) was extracted and analyzed using HPLC.

### 2.3 Cell culture

A7r5 VSMCs were obtained from Bioresource Collection and Research Center (BCRC; Hsinchu, Taiwan) and cultured in Dulbecco’s modified Eagle’s medium (DMEM; Life Technologies; Carlsbad, CA, USA) supplemented with 10% fetal bovine serum (FBS) and a 1% antibiotic/antimycotic solution at 37°C in an incubator containing 5% CO2. Before conducting our experiments, we precultured VSMCs in serum-free medium for 24 h. U937 cells were obtained from the BCRC and grown in Roswell Park Memorial Institute (RPMI)-1640 medium (Life Technologies) supplemented with 10% FBS and 1% antibiotic/antimycotic solution at 37°C in an incubator containing 5% CO2. Human EPCs were obtained from CELLvo™ Human Endothelial Progenitor Cells (StemBioSys, San Antonio, TX) and then cultured in EGM-2 medium. The cells were then plated in human fibronectin-coated tissue culture chambers and cultured at 37°C in a humidified atmosphere containing 5% CO2. The medium was changed every 3 days. All cell lines were used for experiments between passages 3 and 10.

### 2.4 In silico analysis

The best fit of HGK in the activating domain of PDK1 was identified using AutoDock Vina.

Cell viability and cytotoxicity assays

VSMCs were treated with HGK, PDGF-BB, or MK2206 for different times, as indicated, and cell viability and cytotoxicity were determined using MTT and lactate dehydrogenase (LDH) assays.

### 2.5 Western blot analysis

Treated cell and tissue lysates were subjected to sodium dodecyl sulfate–polyacrylamide gel electrophoresis (SDS–PAGE) for separation and were then transferred to polyvinylidene fluoride (PVDF) membranes (Millipore, Bedford, MA, USA), which were incubated with the appropriate primary antibodies (all diluted 1:1000 in 1.5% bovine serum albumin (BSA)) overnight at 4°C before an incubation with the appropriate HRP-conjugated secondary antibodies (all diluted 1:6000) for 1 h at room temperature. Immunoreactivity was detected with enhanced chemiluminescence and quantified using Gel-Pro software. The intensities of the target protein bands were normalized to those of β–actin, GAPDH, α-tubulin or Na+/K+-ATPase.

### 2.6 Immunofluorescence staining

VSMCs plated on cover glasses were pretreated with HGK or MK2206 for 1 h and were then treated with PDGF-BB for the indicated times. After treatment, the cells were fixed with 4% paraformaldehyde for 30 min and then permeabilized with 0.05% Triton X-100 for 2 min. The treated cells were then blocked with 10% normal horse serum for 1 h and incubated with primary antibodies (all diluted 1:100 in blocking solution) at 4°C overnight. Next, the cells were incubated with FITC- or TRITC-conjugated secondary antibodies (all diluted 1:200 in blocking solution) for 1 h at room temperature. The cells were counterstained with DAPI and examined using a fluorescence microscope.

### 2.7 Cell cycle analysis

VSMCs were treated first with HGK or MK2206 for 1 h and then with 20 ng/ml PDGF-BB for another 24 h. The treated cells were trypsinized and stained with PI. Cell cycle progression was assessed with a FACScan flow cytometer, and data were analyzed with ModFit software.

Smooth muscle cell wound healing assay and phalloidin staining

After VSMCs were treated with HGK or MK2206 for 1 h, wounds were created in the cell monolayers by scratching with a sterile pipette tip, creating 250-μm cell-free paths. Then, the medium was replaced with fresh medium to remove cell debris. The cells were rinsed and cultured in complete medium containing vehicle control or 20 ng/ml PDGF-BB with or without 10 μM HGK, 1 μM MK2206, 1 μM BX795 or 5 nM PS48. Cell migration in the wounded area was evaluated after 24 h. Time-lapse images were acquired at 10-min intervals throughout a 24-h period after wounding. The cell migration rate was measured after the acquisition of sequential time-lapse images. Analyses were performed on sequential phase contrast images with MetaMorph software. The number of lamellipodia formed was determined at 9 h after wounding. The treated cells were fixed and then stained with phalloidin and DAPI. Cells in six fields were counted under a 20× objective to determine the number of lamellipodia formed.

### 2.8 BrdU incorporation assay

VSMCs were cultured on gelatin-coated coverslips and serum-starved for 24 h. After pretreatment with 10 μM HGK or 1 μM MK2206 for 1 h, the cells were treated with 20 ng/ml PDGF-BB and 40 mg/ml BrdU for another 24 h. The cells were then fixed with a 95% ethanol/5% acetic acid solution for 30 min and treated with 1 N hydrochloric acid (HCl) for 10 min. Subsequently, the cells were treated with sodium borate to neutralize the HCl. The cells were subsequently incubated first with an anti-BrdU antibody or normal IgG overnight at 4°C and then with TRITC-conjugated goat anti-mouse IgG. The cells were then counterstained with 1 μg/ml DAPI and observed under a fluorescence microscope. Cells in six fields were counted under a 20× objective to determine the number of BrdU-positive nuclei and the total number of nuclei (DAPI-positive nuclei).

### 2.9 Adhesion assay

Cultured U937 cells were stained with BCECF-AM. Stained U937 cells were added to treated cells and incubated for 60 min at 37°C. Then, the cells were washed and observed under a fluorescence microscope.

### 2.10 Mice femoral artery wire injury model

Male C57BL/6 mice (8 w old) were purchased from the National Laboratory Animal Center. Mice were maintained on a 12-h light–dark cycle with food and water available ad libitum. This protocol was approved by the National Taiwan University College of Medicine and the College of Public Health’s Institutional Animal Care and Use Committee (IACUC 20150293). Mice were anesthetized by administering an intraperitoneal (i.p.) injection of pentobarbital (50 mg/kg). Then, endothelial denudation (ED) of the femoral artery was conducted using a spring wire (0.38-mm diameter, No. C-SF-15-15, Cook, Bloomington, IN, USA). Acetaminophen (2.5 mg/kg) was administered in drinking water to provide analgesia. The mice were divided into 2 groups in a completely randomized design: (1) group I (ED/control), in which vehicle (dimethyl sulfoxide, DMSO) was administered by daily intraperitoneal injections for 4 weeks; and (2) group II (ED/HGK), in which HGK was administered (1 mg/kg/day in DMSO) by daily intraperitoneal injections for 4 weeks. Before and 5 days after transluminal mechanical injury, peripheral blood was collected to determine the number of circulating EPCs. Twenty-eight days after transluminal mechanical injury, all mice were euthanized by administering isoflurane anesthesia followed by cervical dislocation. The injured femoral arteries were gently dissected, fixed with 4% paraformaldehyde, embedded in optimal cutting temperature (OCT) compound, and cross-sectioned for a morphometric analysis and immunohistochemistry. Every tenth femoral artery section was collected and stained with resorcin–fuchsin solution (Sigma) and Masson’s trichrome (Scytek Laboratories) to analyze neointima formation and the collagen content. The mean intima/media cross-sectional area (I/M) ratio was calculated using Image-Pro Plus 4.5 software and used to indicate neointima formation or restenosis. The mean collagen and elastin contents were determined using ImageJ software.

### 2.11 Transwell assay

EPCs were treated with or without 1 μM MK2206 for 1 h, seeded in the upper compartment, and incubated with or without 10 μM HGK in the lower compartment for another 6 h. The number of transmigrated cells was determined after 6 h by performing Coomassie blue staining and counting.

### 2.12 RNA sequencing

Total RNA was extracted from treated VSMCs using TRIzol reagent (Invitrogen, Carlsbad, CA, USA), and the extracted RNA samples were sent to Allbio (Taipei, Taiwan) for analysis. Briefly, mRNA was reverse transcribed to cDNAs. Poly(dA) tails were added to the cDNAs in a single reaction, and multiple adaptors were then ligated to both ends of the double-stranded cDNAs. The library was validated and quantified with a Qsep100 automated DNA analyzer (Bioptic, Taiwan, China) and a Qubit 3.0 fluorometer (Invitrogen). Libraries were sequenced with the HiSeq platform.

### 2.13 Evans blue staining

Seven days after transluminal mechanical injury, 50 μl of Evans blue (Sigma) (10%) were administered intravenously before euthanasia. After fifteen minutes, the mice were perfused, and the injured femoral arteries were gently dissected to evaluate the denuded vessel area. The ratio of the Evans blue-stained area to the total femoral artery area was calculated.

### 2.14 Immunoprecipitation

VSMCs were pretreated with HGK or MK2206 for 1 h and then treated with PDGF-BB for another 6 min. The treated cell lysates (400 μg) were incubated with 4 μg of an antibody specific for the Rac family small GTPase 1 (Rac1) at 4°C overnight. Then, the cell lysates were incubated with 50 μl of Protein A agarose beads (Sino Biological, Chesterbrook, PA, USA) at 4°C for another 2 h. Afterward, the beads were washed three times with lysis buffer, and the bound proteins were then eluted by heating (95°C, 10 min) and analyzed using SDS–PAGE and Western blotting. The membrane was incubated first with the appropriate anti-Rac1 and anti-ATP7A antibodies overnight at 4°C and then with the appropriate HRP-conjugated secondary antibodies for 1 h at room temperature. Immunoreactivity was detected with enhanced chemiluminescence and quantified using Gel-Pro software.

### 2.15 Statistical analyses

Statistical analyses were conducted only for studies with a group size of at least 5. All values are provided as the means ± standard deviations (SDs). Statistical comparisons were performed using two-tailed Student’s t test and one-way analysis of variance (ANOVA) followed by Tukey’s post hoc test. Significance was defined as a p value<0.05.

## 3. Results

### 3.1 Isolation and identification of HGK

1H-NMR (400 MHz, DMSO-d6): δ 12.98 (1H, s, D2O exchangeable, OH-5), 9.44 (2H, br.s, D2O exchangeable, OH), 7.46 (1H, dd, J = 8.0, 2.4 Hz, H-6’), 7.44 (1H, d, J = 2.4, H-2’), 6.91 (1H, d, J = 8.0 Hz, H-5’), 6.73 (1H, s, H-3), 6.72 (1H, d, J = 2.4 Hz, H-8), 6.38 (1H, d, J = 2.4 Hz, H-6), 3.88 (3H, s, OCH3) (Figure 1A); 13C-NMR (100 MHz, DMSO-d6): δ 182.3 (C-4), 165.6 (C-7), 164.8 (C-2), 161.7 (C-5), 157.7 (C-9), 150.3 (C-4’), 146.3 (C-3’) 121.9 (C-1’), 119.6 (C-6’), 116.5 (C-5’), 114.0 (C-2’), 105.2 (C-10), 103.6 (C-3), 98.4 (C-6), 93.1 (C-8), 56.5 (OCH3) (Figure 1B). The molecular weight of HGK was recognized based on the Fast atom bombardment mass spectrometry (FAB-MS) peak at m/z 301 ([M+H]+, 5%) (Figure 1C). These data indicated the structure of HGK (Figure 1D).

**Figure 1.**
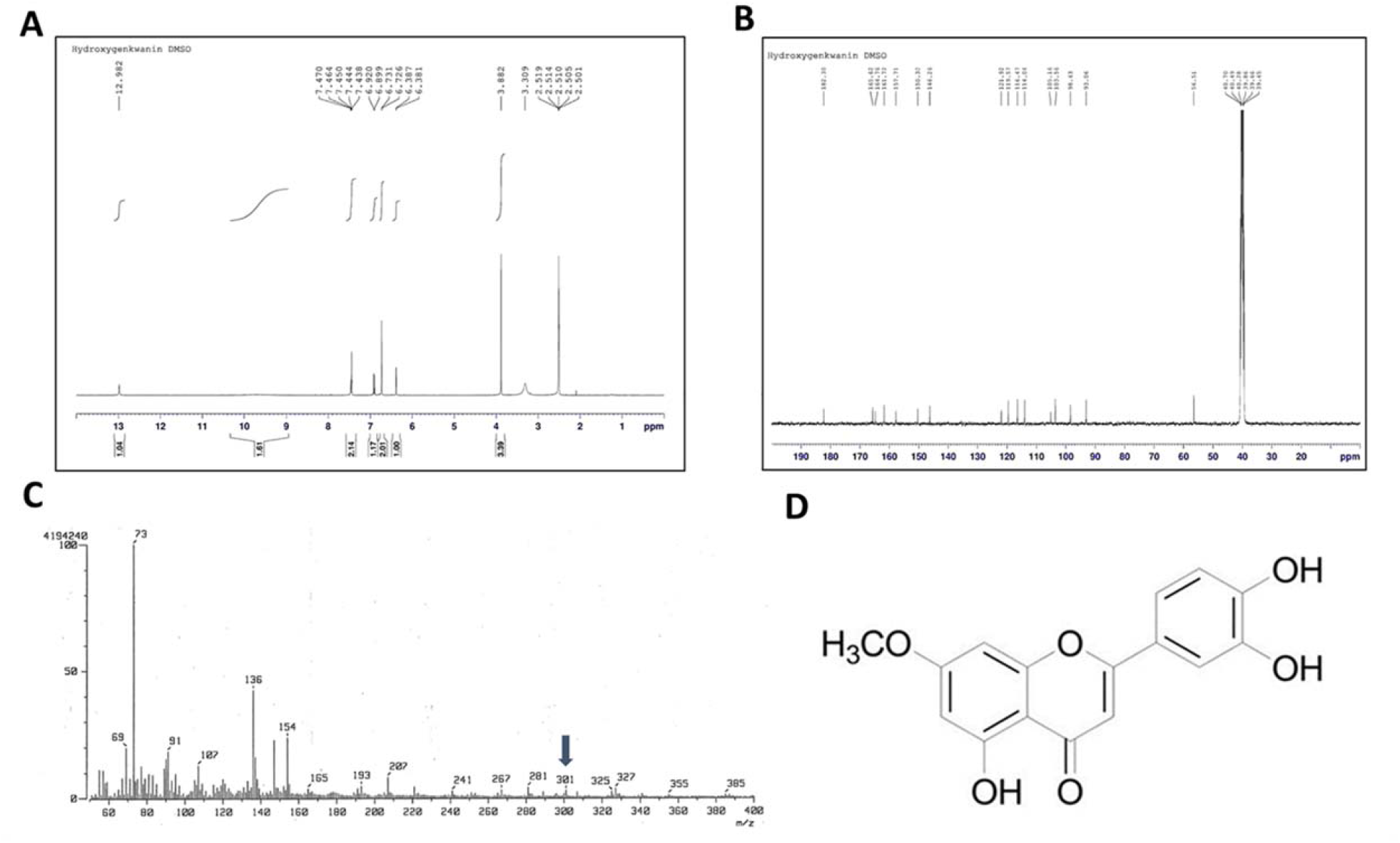
Identification of the HGK structure. (A) 1H NMR, (B) 13C NMR, and (C) FAB-MS analyses of HGK. (D) Chemical structure of HGK.

### 3.2 HGK reduces platelet-derived growth factor-BB (PDGF-BB)-induced VSMC proliferation and migration by inhibiting the AKT pathway

Abnormal VSMC proliferation and migration play important roles in neointimal hyperplasia and restenosis. After chemokine stimulation, contractile VSMCs transform into synthetic VSMCs. PDGF, a potent growth factor present in the injured vessel wall, contributes to VSMC phenotypic switching and VSMC proliferation and migration involved in vascular remodeling by activating multiple intracellular signal transduction pathways. Activated synthetic VSMCs exhibit increased proliferation, migration and extracellular matrix production (3). The matrix metallopeptidase 9 (MMP-9) expression level is an indicator of restenosis (13). Western blot analysis showed that compared with treatment with PDGF-BB alone, combined treatment with 10 μM HGK significantly reduced PDGF-BB-induced expression of synthetic VSMC markers (MMP-9 and Connexin-43 (Cx43)) and increased expression of VSMC contractile marker α-smooth muscle actin (α-SMA) (Figure 2A). MTT, LDH 5-bromo-2’-deoxyuridine (BrdU) incorporation assay, flow cytometry and western blotting analysis all showed that 10 μM HGK pre-treatment significantly reduced PDGF-BB-induced proliferation, BrdU incorporation, the proportion of S-phase cells and expression of proliferation-regulating proteins (Cyclin D1, CDK4, Cyclin E and CDK2) without inducing cell death and increased P27 expression (Figure 2B-D, Supplemental Figure 1A-B). In addition to inhibiting proliferation, HGK also reduced PDGF-BB-induced VSMC migration and the formation of lamellipodia at the leading edge (Figure 2E, Supplemental Figure 1C). Accumulating evidence has shown that AKT activation plays an important role in VSMC proliferation and migration (14). In the present study, compared to PDGF-BB treatment alone, combined treatment with HGK markedly reduced PDGF-BB-induced AKT phosphorylation (Figure 2F). In addition, treatment with HGK reduced PDGF-BB-induced VSMC proliferation, the expression of proliferation-regulating proteins (Cyclin D1, CDK4, Cyclin E and CDK2), the expression of MMP-9, migration, and lamellipodia formation and increased the expression level of P27. These effects of HGK on PDGF-BB-treated VSMCs were similar to those of MK2206 (an AKT inhibitor) (Figure 2G-J, Supplemental Figure 2A-C). Based on these data, HGK reduced PDGF-BB-induced VSMC proliferation and migration by regulating AKT.

**Figure 2.**
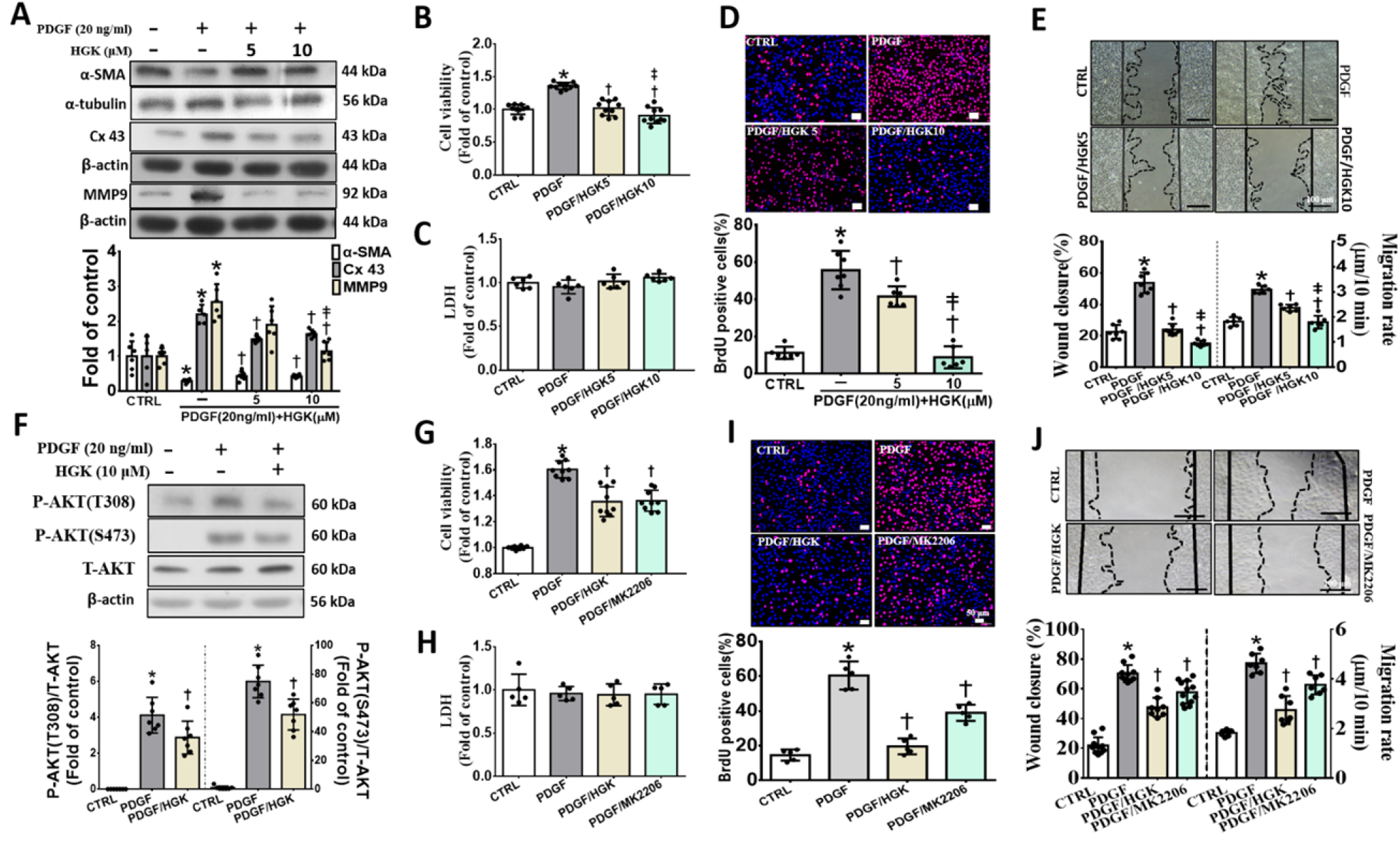
HGK decreases PDGF-BB-induced VSMC proliferation and migration by inhibiting the AKT pathway. Serum-starved VSMCs were pretreated with 5 or 10 μM HGK for 1 h and then treated with 20 ng/ml PDGF-BB (PDGF) for another 24 h. (A) Western blot analyses showed that HGK increased contractile VSMC marker expression and reduced synthetic VSMC marker expression. (B-D) MTT, LDH, and BrdU incorporation assays were performed. These results showed that HGK decreased PDGF-induced VSMC proliferation without inducing cell death. (E) Wound healing assays indicated that HGK decreased PDGF-induced VSMC migration. The black and dotted lines indicate the wound edge on day 0 and day 1, respectively. (F) Western blot analyses showed that HGK reduced PDGF-induced AKT phosphorylation. (B-E) MTT, LDH, BrdU incorporation assays and wound healing assays showed that HGK decreased PDGF-induced VSMC proliferation and migration by inhibiting AKT phosphorylation. The black and dotted lines indicate the wound edge on day 0 and day 1, respectively. The scale bar represents 50 μm in (D and I) and 100 μm in (E and J). Nuclei were stained with DAPI. The values are presented as the means ± SDs. *p < 0.05 compared with the CTRL group. †p < 0.05 compared with the PDGF group. ‡p < 0.05 compared with the PDGF/HGK5 group. N=5-7. CTRL, control group. Statistical comparisons were performed using one-way ANOVA.

### 3.3 HGK decreases PDGF-BB-induced VSMC migration by regulating AKT/ATP7A/Rac1 complex formation

Next-generation sequencing (NGS) showed that HGK treatment resulted in characteristics that differed from those induced by PDGF-BB treatment (Figure 3A). The 13 differentially regulated mRNA transcripts were visualized in a volcano plot. A copper-transporting ATPase (ATP7) was one of the most upregulated annotated genes and is indicated with a circle (Figure 3B). A previous study showed that ATP7A plays an important role in PDGF-BB-induced VSMC migration (15, 16). After VSMCs were stimulated with PDGF-BB, ATP7A colocalized with Rac1 and then translocated from the cytosol to the leading edge, where it mediated lamellipodia formation (15). We then investigated the role of the ATP7A/Rac1 complex in the HGK-mediated repression of PDGF-BB-induced VSMC migration. Both the immunoprecipitation and Western blot analyses showed that HGK reduced the level of ATP7A/Rac1 complex formation and inhibited the PDGF-BB-induced translocation of Rac1 from the cytosol to the membrane; in addition, these inhibitory effects were similar to those of MK2206 (an AKT inhibitor). (Figure 3C-D). Similar results were obtained using immunofluorescence staining (Figure 3E). Thus, HGK reduces PDGF-BB-induced VSMC proliferation and migration by inhibiting ATP7A/Rac1 complex translocation through the regulation of the AKT pathway.

**Figure 3.**
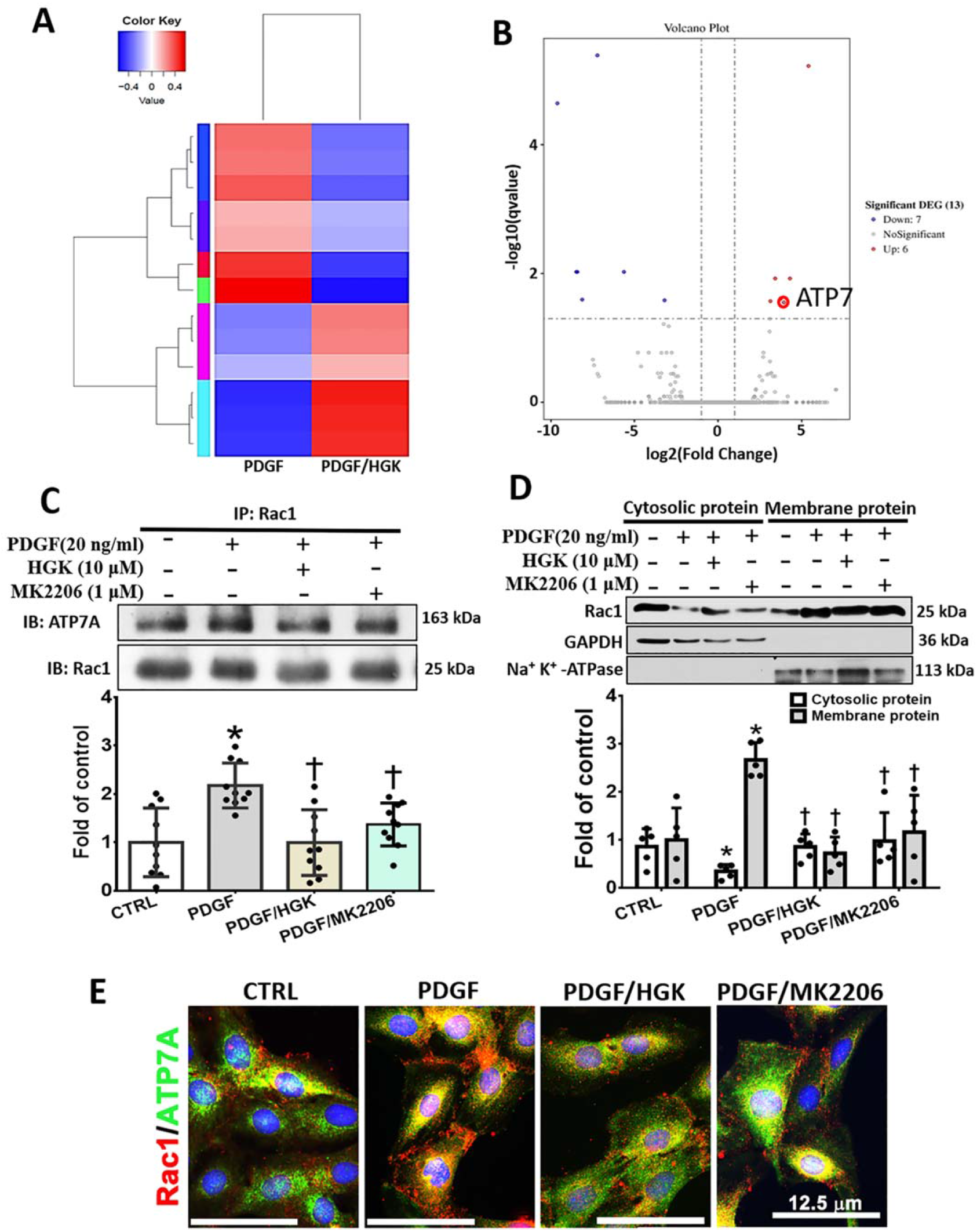
HGK decreases PDGF-BB-induced VSMC proliferation and migration by inhibiting the translocation of the ATP7A/Rac1 complex through the regulation of the AKT pathway. (A) An RNA transcriptome analysis was performed to analyze the gene expression profile of PDGF-BB (PDGF)-treated VSMCs treated with or without HGK(PDGF/HGK). (B) A volcano plot of the 13 differentially regulated mRNA transcripts (shown in blue and red). ATP7 was one of the most upregulated annotated genes and is indicated with a circle. (C) Immunoprecipitation showed that treatment with 10 μM HGK or 1 μM MK2206 decreased ATP7A and Rac1 colocalization. (D) Western blot analysis showed that treatment with 10 μM HGK or 1 μM MK2206 decreased PDGF-induced Rac1 translocation from the cytosolic compartment to the membrane compartment. GAPDH or Na+/K+-ATPase was processed in parallel as an internal control for protein loading. (E) Immunofluorescence staining showed that pretreatment with 10 μM HGK or 1 μM MK2206 decreased PDGF (20 ng/ml)-induced ATP7A and Rac1 colocalization and translocation. The scale bar represents 12.5 μm in (E). Nuclei were stained with DAPI. The values are presented as the means ± SDs. *p < 0.05 compared with the CTRL group. †p < 0.05 compared with the PDGF group. N=5-10. CTRL, control group. Statistical comparisons were performed using one-way ANOVA.

### 3.4 HGK reduces tumor necrosis factor-α (TNF-α)-induced VSMC inflammation and thrombosis by inhibiting the AKT pathway

Endothelial dysfunction and injury trigger the infiltration and adhesion of inflammatory leukocytes and induce excessive fibrin deposition to cause thrombosis and restenosis (17). Numerous studies have shown that TNF-α production increases and aggravates restenosis progression after angioplasty or vessel injury (18). We evaluated the inhibitory effects of HGK on inflammation and thrombosis in TNF-α-treated VSMCs and found that a high concentration of HGK significantly inhibited the TNF-α-induced expression of inflammatory adhesion molecules (such as intercellular adhesion protein 1 (ICAM-1), E-selectin and vascular cell adhesion protein 1 (VCAM-1)) and thromboinflammatory factors (TF and plasminogen activator inhibitor 1 (PAI-1)) compared with that in the TNF-α treatment group (Figure 4A-B). We further investigated the molecular mechanisms underlying the inhibitory effects of HGK on TNF-α induced inflammation and thrombosis. The AKT pathway plays a key role in inflammation and thrombosis (19). Our data showed that HGK markedly inhibited TNF-α-induced AKT phosphorylation (Figure 4C). The inhibitory effects of HGK on the TNF-α-induced expression of VSMC inflammatory adhesion molecules (ICAM-1 and E-selectin) and thromboinflammatory factors were similar to those of MK2206 (Figure 4D-E). In addition, the monocyte adhesion assay showed that compared with TNF-α treatment alone, HGK pretreatment significantly reduced the increase in monocyte adhesion to TNF-α-treated VSMCs. The involvement of ICAM-1, E-selectin and VCAM-1 in the adhesion of monocytes to TNF-α-treated VSMCs was examined by pretreating cells with 1 μg/ml anti-VCAM-1, anti-E-selectin and anti-VCAM-1 antibodies for 1 h before an incubation with TNF-α for another 24 h. Pretreatment with antibodies specific for adhesion molecules resulted in a significant reduction in the adhesion of monocytes to TNF-α-treated VSMCs compared to that in the groups treated with TNF-α and cotreated with TNF-α and the IgG isotype control. These results indicated that ICAM-1, E-selectin and VCAM-1 play major roles in monocyte adhesion to TNF-α-treated VSMCs. The adhesion of monocytes to TNF-α-treated VSMCs was also inhibited by 1 μM MK2206 (Figure 4F), and these inhibitory effects were similar to those of pretreatment with antibodies specific for inflammatory adhesion molecules or HGK. Therefore, HGK inhibits both monocyte adhesion to TNF-α-stimulated VSMCs and thrombosis by inhibiting ICAM-1, E-selectin, TF and PAI-1 expression and this effect may be partially mediated by regulating the AKT pathway.

**Figure 4.**
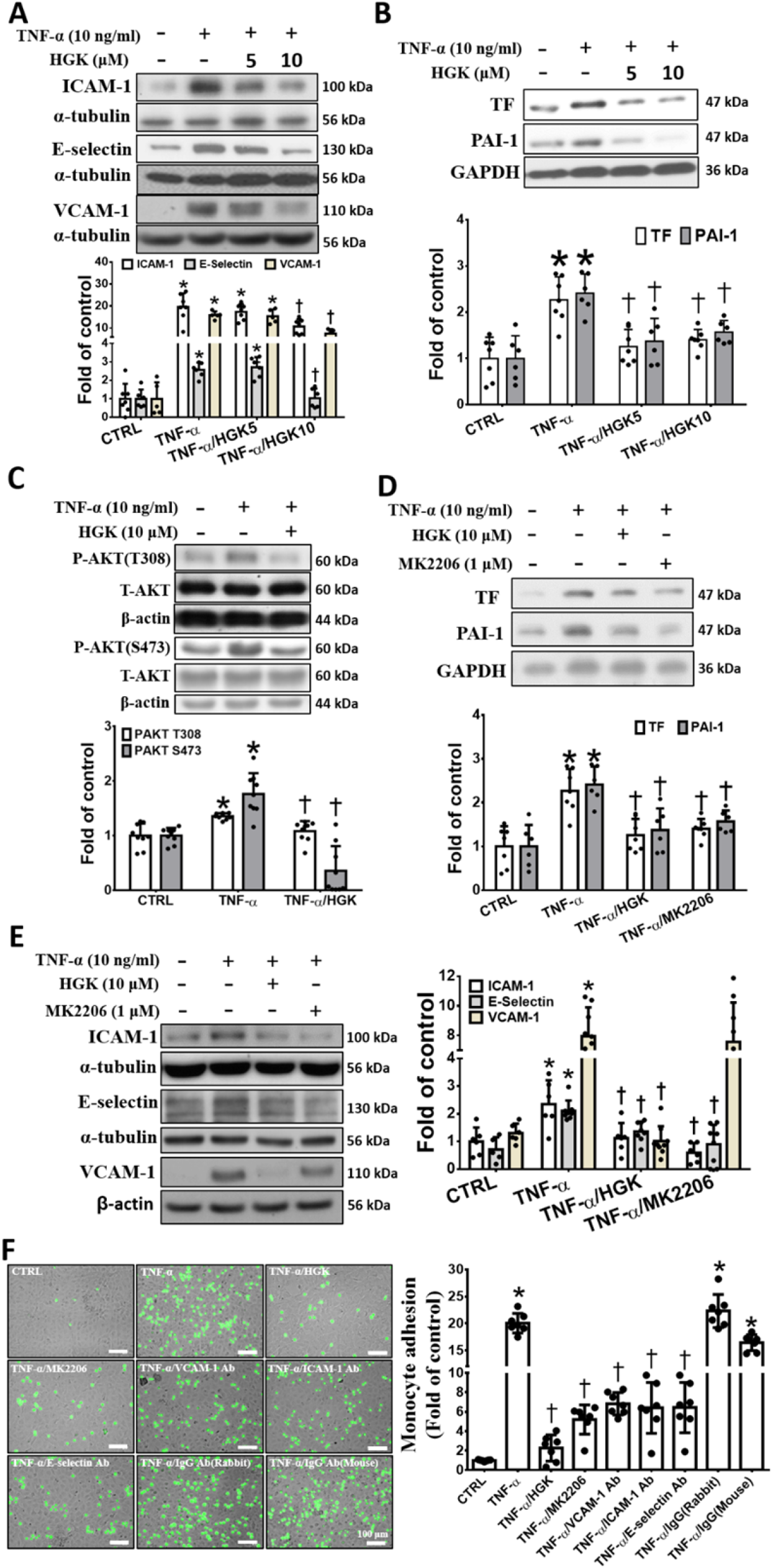
HGK decreases TNF-α-induced thrombosis and inflammation by inhibiting the AKT pathway in VSMCs. Serum-starved VSMCs were pretreated with 5 or 10 μM HGK for 1 h and then treated with 10 ng/ml TNF-α for different times. (A-B) Western blot analysis showed that HGK decreased the expression levels of adhesion molecules, TF and PAI-1 in TNF-α-treated VSMCs. (C) Western blot analyses showed that HGK reduced TNF-α-induced AKT phosphorylation. (D-E) Serum-starved VSMCs were pretreated with 5 or 10 μM HGK or 1 μM MK2206 for 1 h and were then treated with 10 ng/ml TNF-α for another 24 h. Western blot analyses showed that treatment with HGK or MK2206 decreased the expression levels of adhesion molecules, TF and PAI-1 in TNF-α-treated VSMCs. (F) Serum-starved VSMCs were pretreated with 10 μM HGK, 1 μM MK2206, and 1 μg/ml anti-VCAM-1, anti-ICAM-1, and anti-E-selectin antibodies or IgG for 1 h. These treated VSMCs were then stimulated with 10 ng/ml TNF-α for another 24 h. BCECF-labeled U937 monocytes (green) were cocultured with the treated VSMCs for 1 h. Adhesion assays showed that the HGK pretreatment decreased TNF-α-induced VSMC inflammation. The scale bar represents 100 μm in (D). The values are presented as the means ± SDs. *p < 0.05 compared with the CTRL group. †p < 0.05 compared with the TNF-α group. N=5-9. CTRL, control group. Statistical comparisons were performed using one-way ANOVA.

### 3.5 HGK reduces PDGF-BB- or TNF-α-induced VSMC proliferation, migration and inflammation by inhibiting the AKT/mammalian target of rapamycin (mTOR)/S6 kinase (S6K) pathway

mTOR-S6K signaling plays important roles in cell growth, proliferation, migration, and inflammation (20). Western blot analyses showed that both HGK and MK2206 reduced the levels of phosphorylated mTOR and S6K in PDGF-BB- and TNF-α-treated VSMCs (Figure 5A-B). The inhibitory effects on PDGF-BB- or TNF-α-treated VSMCs were similar between the HGK and MK2206 treatment groups. These results indicated that HGK inhibits VSMC proliferation, migration and inflammation by suppressing mTOR-S6K signaling through the AKT pathway.

**Figure 5.**
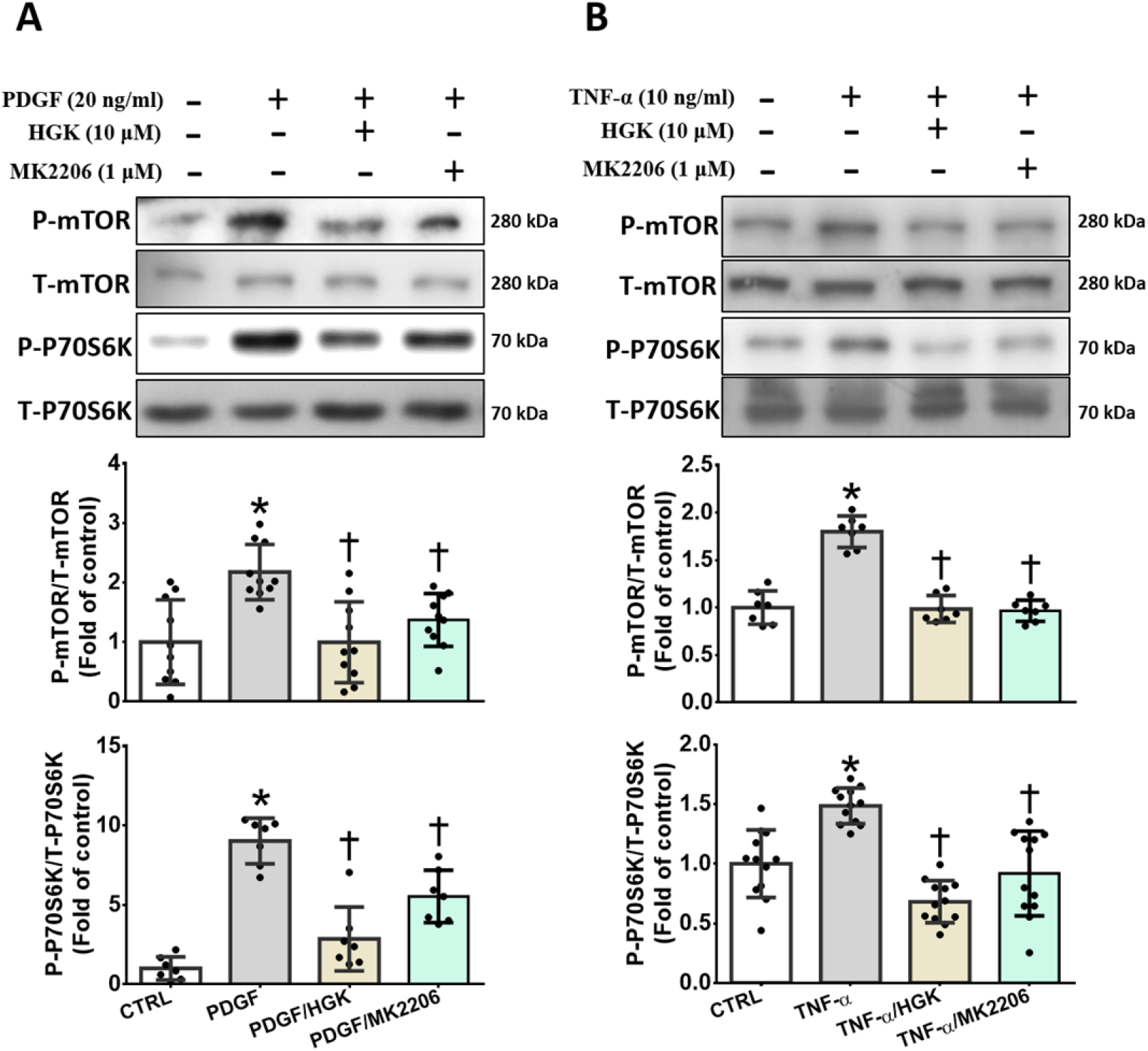
HGK decreases PDGF-BB- or TNF-α-induced mTOR/S6K phosphorylation in VSMCs by inhibiting AKT activation. Serum-starved VSMCs were pretreated with 10 μM HGK or 1 μM MK2206 for 1 h and then treated with 20 ng/ml PDGF-BB (PDGF) or 10 ng/ml TNF-α for another 24 h. (A-B) Western blot analyses showed that HGK or MK2206 pretreatment decreased PDGF- and TNF-α-induced mTOR and S6K phosphorylation. The values are presented as the means ± SDs. *p < 0.05 compared with the CTRL group. †p < 0.05 compared with the PDGF or TNF-α group. N=7-12. CTRL, control group. Statistical comparisons were performed using one-way ANOVA.

We further investigated the molecular mechanisms underlying the inhibitory effects of HGK on VSMC proliferation, migration and inflammation. PDK1 has been reported to play an important role in regulating proliferation, migration and inflammation by activating AKT via phosphorylation at threonine 308 (Thr308) and serine 473 (Ser473) (21). The docking analysis showed that HGK has a strong binding affinity for PDK1 (Figure 6A). In addition, HGK treatment decreased PDK1 phosphorylation in PDGF-BB- and TNF-α-treated VSMCs (Figure 6B). Moreover, PDGF-BB- or TNF-α-induced phosphorylation of AKT at Ser473 and Thr308 was also inhibited by the HGK pretreatment. Similar inhibitory effects were observed on the group pretreated with BX795 (a PDK1 inhibitor) (Figure 6C-D). Based on these results, HGK decreases PDGF-BB- or TNF-α-induced AKT phosphorylation in VSMCs through the PDK1 pathway.

**Figure 6.**
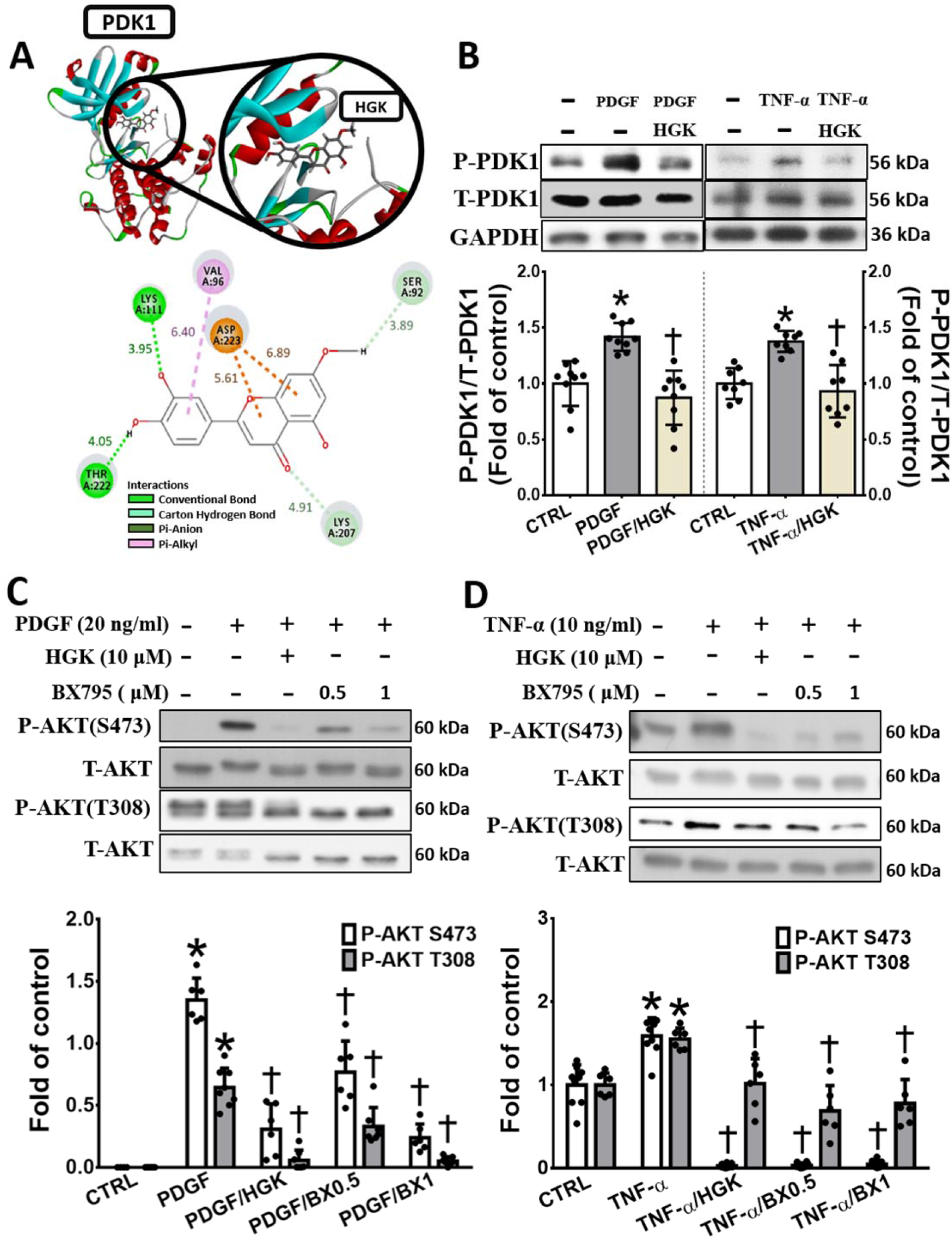
HGK decreases PDGF-BB- or TNF-α-induced AKT phosphorylation in VSMCs by inhibiting PDK1. (A) The in silico analysis showed that HGK has a high binding affinity for PDK1. (B) Western blot analyses showed that HGK decreased PDGF-BB (PDGF)- or TNF-α-induced PDK1 phosphorylation. (C) Serum-starved VSMCs were pretreated with 10 μM HGK or 0.5-1 μM BX795 (a PDK1 inhibitor) for 1 h and were then treated with 20 ng/ml PDGF-BB or 10 ng/ml TNF-α for another 15 or 10 min, respectively. Western blot analyses showed that HGK and BX795 decreased PDGF-BB- or TNF-α-induced AKT phosphorylation at Ser473 (S473) and Thr308 (T308). The values are presented as the means ± SDs. *p < 0.05 compared with the CTRL group. †p < 0.05 compared with the PDGF or TNF-α group. N=6-9. CTRL, control group. Statistical comparisons were performed using one-way ANOVA.

Cell viability and wound healing assays were performed to further elucidate the role of PDK1 in the HGK-mediated inhibition of PDGF-BB-induced VSMC proliferation and migration. The inhibitory effects of HGK on PDGF-BB-treated VSMC proliferation and migration were similar to those of the BX795 pretreatment and HGK/BX795 cotreatment. In addition, HGK inhibited the proliferation and migration of VSMCs treated with PS48 (a PDK1 activator). (Figure 7A-B). The inhibitory effects of HGK treatment, BX795 treatment or HGK/BX795 cotreatment on TNF-α-induced expression of inflammatory adhesion molecules and thromboinflammatory factors in VSMCs were similar, but TNF-α-induced VCAM-1 expression was not suppressed by the BX795 pretreatment. HGK also suppressed the expression of inflammatory adhesion molecules and thromboinflammatory factors in PS48-treated VSMCs. However, VCAM-1 expression in TNF-α-treated VSMCs was not inhibited by BX795 treatment and was induced by PS48 treatment (Figure 7C-F). Moreover, the Western blot analysis showed that pretreatment with either HGK or BX795 reduced the levels of phosphorylated mTOR and S6K in PDGF-BB- and TNF-α-treated VSMCs (Figure 8A-B) to a similar degree. Furthermore, compared to the control treatment, PS48 treatment significantly induced the phosphorylation of mTOR and S6K, and compared to PS48 treatment, HGK treatment substantially reduced the levels of mTOR and S6K phosphorylation (Figure 8C). Therefore, HGK inhibits VSMC proliferation, migration and inflammation mainly by inhibiting PDK1 activation and subsequently regulating the AKT/mTOR/S6K pathway.

**Figure 7.**
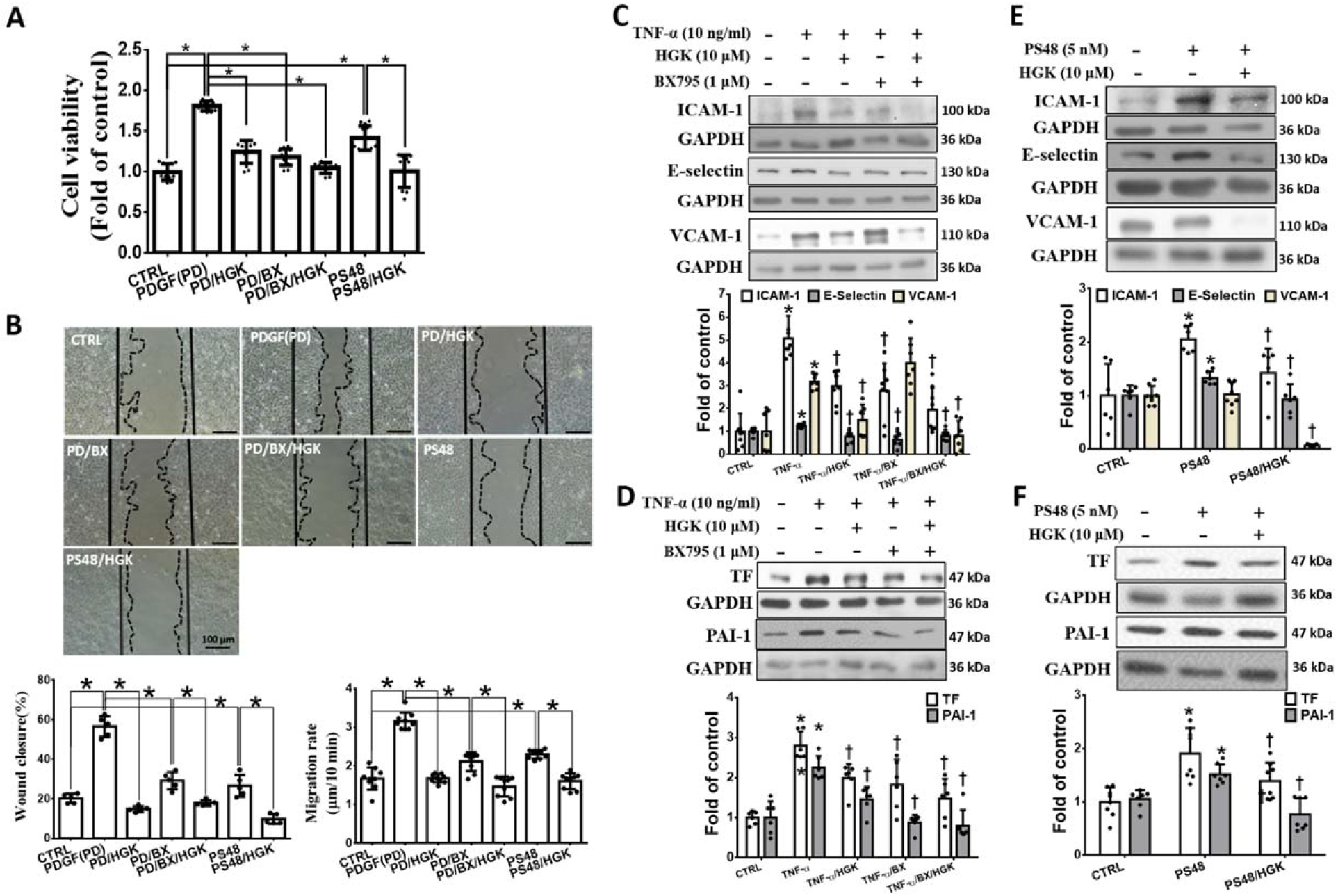
HGK decreases PDGF-BB- or TNF-α-induced VSMC proliferation and migration through PDK1 inhibition. Serum-starved VSMCs were pretreated with 10 μM HGK or 1 μM BX795 (a PDK1 inhibitor) for 1 h and were then treated with 20 ng/ml PDGF-BB (PDGF), 10 ng/ml TNF-α or 5 nM PS48 (a PDK1 activator) for another 24 h. (A-B) MTT and wound healing assays demonstrated that HGK decreased PDGF (PD)- or PS48-induced VSMC proliferation and migration through PDK1 inhibition. The black and dotted lines indicate the wound edge on day 0 and day 1, respectively. *p <0.05. The scale bar represents 100 μm in (B). (C-F) Western blot analysis showed that HGK decreased TNF-α- or PS48-induced expression of adhesion molecules, TF and PAI-1 in VSMCs through PDK1 regulation. The values are presented as the means ± SDs. *p <0.05 vs. the CTRL group. †p <0.05 vs. the PDGF group. N=5-15. CTRL, control group. Statistical comparisons were performed using one-way ANOVA.

**Fig. 8.**
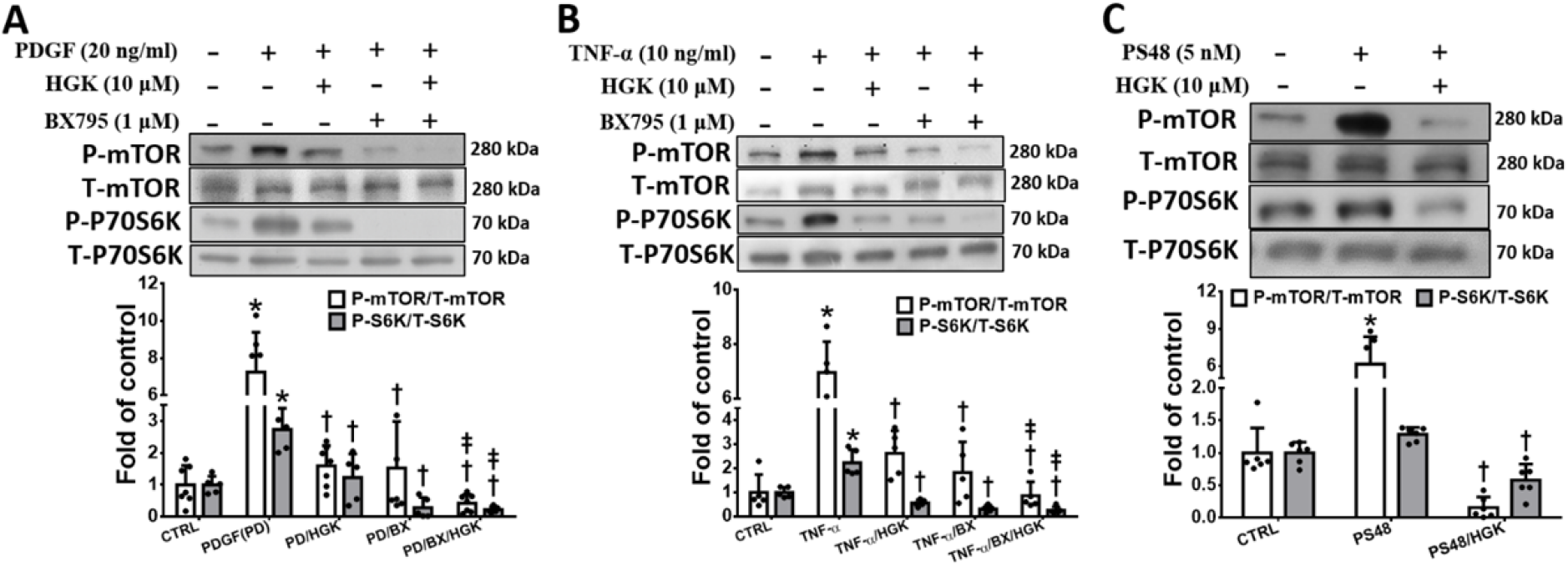
HGK decreases the phosphorylation of mTOR and S6K in PDGF-BB- or TNF-α-induced VSMCs through PDK1 inhibition. Serum-starved VSMCs were pretreated with 10 μM HGK or 1 μM BX795 (a PDK1 inhibitor) for 1 h and were then treated with 20 ng/ml PDGF-BB (PDGF), 10 ng/ml TNF-α- or 5 nM PS48 (a PDK1 activator) for another 24 h. (A-C) Western blot analysis showed that HGK decreased PDGF-, TNF-α- or PS48-induced mTOR and S6K phosphorylation through PDK1 inhibition. The values are presented as the means ± SDs. *p <0.05 vs. the CTRL group. †p <0.05 vs. the TNF-α or PS48 group. ‡p <0.05 vs. the PDGF/HGK or TNF-α/HGK group. N=5-8. CTRL, control group. Statistical comparisons were performed using one-way ANOVA.

### 3.6 HGK enhances the chemotaxis of circulating endothelial progenitor cells (EPCs) and reendothelialization after mouse femoral artery denudation

Homing of circulating EPCs and reendothelization inhibit neointimal hyperplasia formation and restenosis progression (22). Peripheral blood was collected 5 days after endothelial denudation to evaluate the therapeutic effects of HGK on the homing of circulating EPCs and reendothelization. The flow cytometry analysis showed that HGK treatment increased the number of circulating EPCs in the blood (Figure 9A). Similar results were observed in the Transwell assay (Figure 9B). These findings showed that HGK treatment increases the chemotaxis of circulating EPCs. In addition, Evans blue staining showed that HGK reduced the degree of vascular injury at 7 days after endothelial denudation (Figure PC). Moreover, CD31 staining revealed that HGK promoted reendothelization in denuded femoral arteries at 28 days after endothelial denudation (Figure 9D). The aforementioned data showed that HGK promotes EPC chemotaxis and reendothelization.

**Fig. 9.**
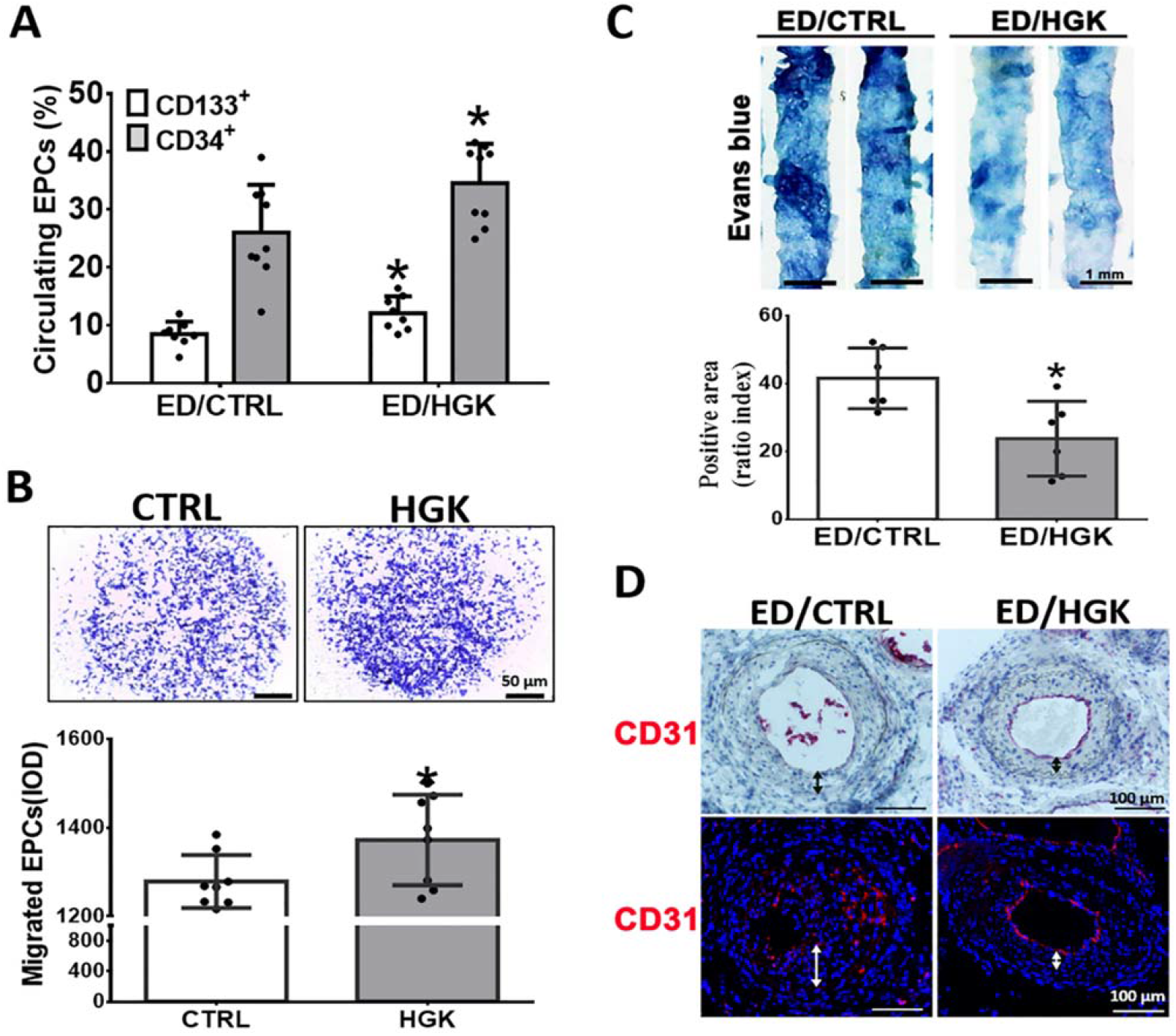
HGK enhances the chemotaxis of circulating EPCs and reendothelialization after femoral artery denudation in mice. (A) The number of circulating blood EPCs was determined by flow cytometry 5 days after endothelial denudation (ED). Treatment with HGK (1 mg/kg) increased the number of circulating EPCs after ED. (B) Transwell assays showed that HGK promoted EPC migration. (C) Denuded femoral arteries were stained with Evans blue dye 7 days after ED. Femoral artery staining demonstrated that HGK promoted reendothelialization. (D) Immunohistochemical staining for CD31 was performed. HGK treatment increased CD31 expression (red) in denuded femoral arteries at 28 days after ED. Nuclei were stained with DAPI or hematoxylin. The thickness of neointimal hyperplasia is indicated by the double-headed arrows. The values are presented as the means ± SDs. *p <0.05 vs. the ED/CTRL or CTRL group. N=5-12. CTRL, control group. Statistical comparisons were performed using Student’s t test and one-way ANOVA.

### 3.7 HGK reduces restenosis in vivo

Endothelial denudation was performed to evaluate the therapeutic effect of HGK on neointimal hyperplasia. Compared to the vehicle treatment, the HGK treatment significantly reduced neointimal hyperplasia, collagen deposition, and the proliferation of VSMCs and macrophages and increased the elastin content in denuded femoral arteries at 28 days after endothelial denudation (Figure 10A-B). MMP-9 is a marker of synthetic VSMCs and a key factor regulating smooth muscle cell proliferation and migration (23). After angioplasty or vessel injury, the activation of inflammation has been shown to play an important role in neointimal growth. Previous studies showed that TNF-α production and the number of infiltrated macrophages increased soon after angioplasty or vessel injury and that these increases were maintained for at least 2 weeks, thereby contributing to arterial restenosis (24). Immunohistochemical staining showed that the HGK treatment reduced MMP-9, macrophage marker and TNF-α expression levels in denuded femoral arteries (Figure 10C). Inflammation, macrophage activation, monocyte adhesion, platelet activation and thrombus formation are also involved in neointimal hyperplasia, restenosis and thrombosis progression, and HGK also reduced the expression levels of adhesion molecules, TF and PAI-1 in denuded femoral arteries (Figure 10D). In addition, the phospho-PDK1 level was markedly reduced in denuded arteries from the HGK treatment group compared with those from the vehicle treatment group (Figure 10E).

**Fig. 10.**
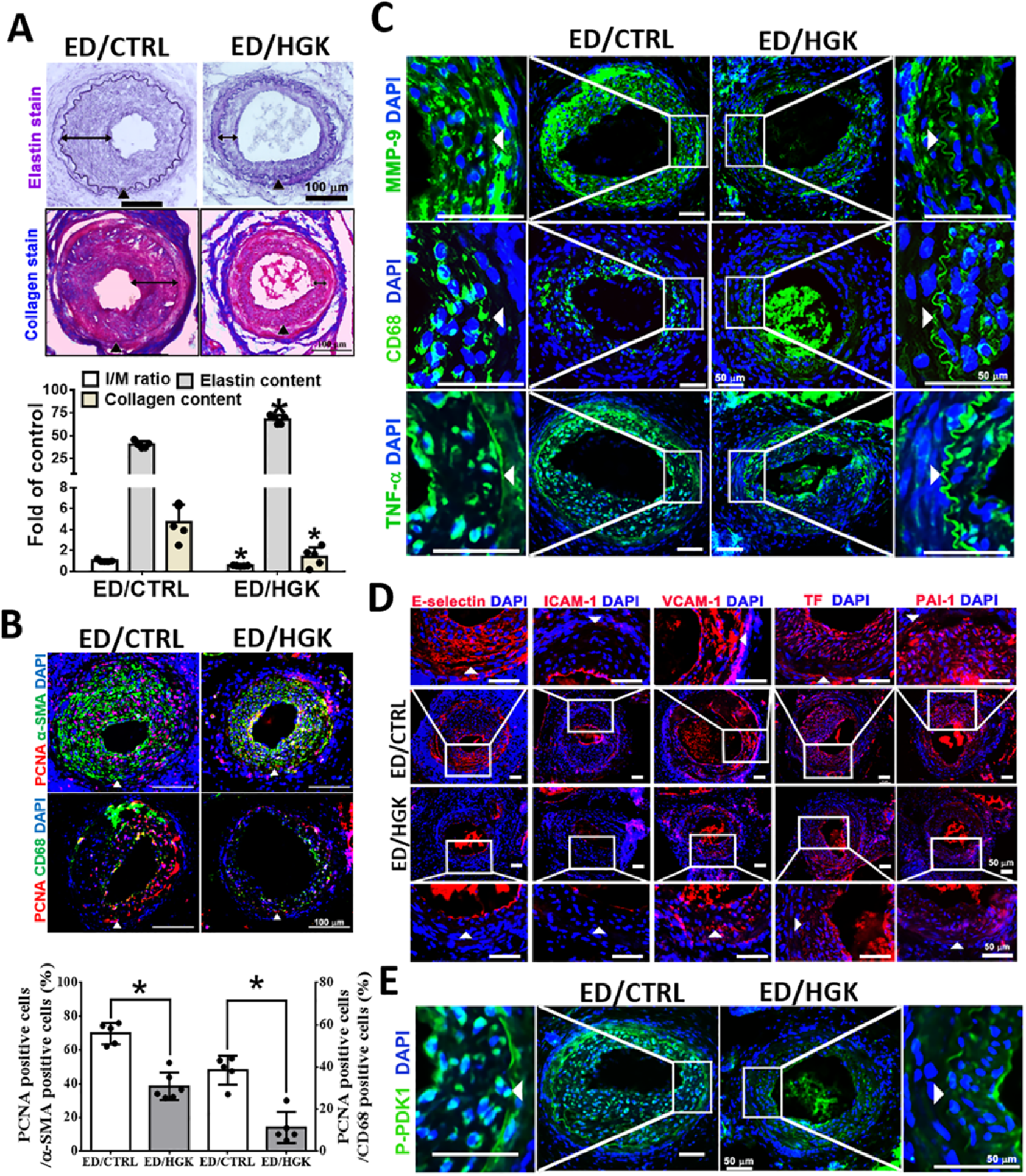
HGK reduces restenosis in vivo. (A)Elastin staining showed that HGK treatment decreased neointimal hyperplasia and increased the elastin content in denuded femoral arteries 28 days after endothelial denudation (ED). Collagen staining showed that HGK treatment decreased collagen expression in denuded femoral arteries 28 days after ED. The black arrowheads indicate the internal elastic lamina. The double-headed arrows indicate the area of neointimal hyperplasia. (B)Immunohistochemical staining for PCNA (red), α-SMA (green) and CD68 (green) was performed. HGK treatment decreased neointimal VSMC and macrophage (CD68) proliferation in denuded femoral arteries 28 days after injury. (C) Immunohistochemical staining showed that HGK treatment decreased the expression levels of MMP-9, a macrophage marker (CD68) and TNF-α in denuded femoral arteries 28 days after ED. (D-E) Immunohistochemical staining showed that HGK treatment decreased the protein levels of adhesion molecules, TF, PAI-1 and phospho-PDK1 in denuded femoral arteries 28 days after ED. The white arrowheads indicate the internal elastic lamina (B-E). The scale bar represents 100 μm in (A-B) and 50 μm in (C-E). Nuclei were stained with DAPI. The values are presented as the means ± SDs. *p <0.05 vs. the ED/CTRL group (control group). N=5. Statistical comparisons were performed using Student’s t test.

## 4. Discussion

The present in vitro studies showed that HGK decreased PDGF-BB-induced VSMC proliferation and migration and TNF-α-stimulated VSMC inflammation mainly by regulating the PDK1/AKT/mTOR/S6K pathway. Furthermore, HGK promoted EPC chemotaxis. In vivo, compared to the vehicle treatment, HGK treatment significantly decreased neointimal hyperplasia by accelerating endothelization; reducing femoral artery denudation-induced intimal SMC proliferation, collagen deposition, MMP-9 production, and inflammatory molecule expression; and increasing the elastin content in the injured artery. These therapeutic effects of HGK may be exerted through PDK1 inhibition.

HGK, a flavonoid extract from the plant Daphne genkwa, exerts potent anti-inflammatory, antitumor, antiproliferative, anti-migratory and immunomodulatory effects with lower toxicity and fewer side effects than synthetic agents (11, 25). In addition to its anticancer activity, HGK also possesses antioxidant activity and may function as a TF inhibitor to prevent thrombosis and subsequent cardiovascular disease progression (12). As shown in the present study, HGK inhibited PDGF-BB-induced VSMC proliferation and migration and reduced TNF-α-stimulated VSMC inflammation. The in silico analysis, in vitro Western blot analysis and functional assays with BX795 and PS48 treatment revealed that HGK may exert its therapeutic effects mainly by regulating the PDK1/AKT/mTOR/S6K pathway. Based on these effects of HGK on PDGF-BB- or TNF-α-induced VSMCs, we found that HGK treatment substantially reduced intimal hyperplasia, VSMC proliferation in the intimal layer, and the expression of MMP-9, adhesion molecules, TF and PAI-1 in the thickened intima in vivo.

Angioplasty is commonly used to treat atherosclerotic plaques or stenosis. Although the occluded vessels were reopened, endothelial denudation or endovascular injury has been observed after angioplasty (26). Endothelial injury promotes platelet aggregation and activation, increases monocyte adhesion and inflammatory cytokine expression, stimulates VSMC proliferation and migration and ultimately results in restenosis (27). Based on accumulating evidence, EPCs play a crucial role in postinjury reendothelialization (28). In addition, emerging evidence has shown that high levels of reactive oxygen species (ROS) in injured vessels are a key feature after vessel denudation (29) and result in dysfunction and migration of endothelial cells and EPCs. Previous studies have shown that HGK exerts antioxidant and free radical scavenging effects. In addition, the results of flow cytometry and CD31 staining showed that HGK treatment increased the number of circulating EPCs on day 5 after femoral artery denudation and accelerated reendothelization in the present study. Furthermore, the results of in vitro assays revealed that HGK increased EPC chemotaxis. Thus, HGK has considerable therapeutic potential for EPC mobilization and endothelial injury repair, mainly through its antioxidant effects.

AKT is a serine/threonine protein kinase with multiple phosphorylation sites and is responsible for multiple behaviors of VSMCs, such as proliferation, migration and differentiation (30). Thr308, which is located in the kinase catalytic region of AKT, is phosphorylated by PDK1, and this phosphorylation is required for AKT activation (31). Then, phospho-AKT (Thr308) phosphorylates AKT at Ser473 via mTORC2 by phosphorylating SIN1 at Thr86 and enhancing mTORC2 kinase activity (32). AKT has a broad range of downstream targets, and accumulating evidence has shown that the AKT/mTOR/S6K signaling pathway is involved in the regulation of several biological processes, such as survival, proliferation, migration and inflammation (33). According to the molecular docking analysis, PDK1 is a major molecular target of HGK. Consistent with this observation, HGK markedly reduced the phosphorylation of PDK1 in PDGF-BB- or TNF-α-treated VSMCs. The increase in AKT (Ser473/Thr308) phosphorylation was substantially inhibited after a PDK1 inhibitor pretreatment, and these effects were similar to those of the HGK pretreatment. The antiproliferative, anti-migratory and anti-inflammatory effects of HGK on PDGF-BB- or TNF-α-stimulated VSMCs were similar to those observed for the BX795 treatment and HGK/BX795 cotreatment. In addition, pretreatment with HGK reduced the proliferation, migration and inflammation of PS48-treated VSMCs. These results indicated that HGK reduced PDGF-BB- or TNF-α-stimulated phosphorylation and activation of AKT mainly by inhibiting PDK1 activation and then substantially reduced VSMC proliferation, migration and inflammation by suppressing AKT activity.

After PDGF-BB or inflammatory cytokine stimulation, contractile VSMCs transform into synthetic VSMCs and then proliferate rapidly and migrate from the tunica media into the intimal layer. The Cu exporter ATP7A (copper-transporting P-type ATPase/Menkes ATPase) plays a key role in PDGF-BB-induced VSMC migration. ATP7A recruits Rac1 and translocates from the trans-Golgi network to the leading edge of VSMCs to promote lamellipodia formation (16). The NGS analysis in the present study indicated differential expression of ATP7 in HGK-treated VSMCs compared with PDGF-BB-treated VSMCs. In addition, the results of immunoprecipitation, Western blot analyses and immunofluorescence staining showed that HGK substantially decreased the PDGF-BB-induced translocation of the ATP7A/Rac1 complex from the cytosol to the cell membrane. In VSMCs, HGK decreased PDGF-BB-induced lamellipodia formation and migration by regulating ATP7A/Rac1 complex formation and localization.

The thromboinflammatory factors TF and PAI-1 are biomarkers for vascular diseases such as atherosclerosis, intimal hyperplasia and restenosis (34, 35). The expression of TF and PAI-1 has been reported to be induced in VSMCs upon stimulation with various cytokines, such as TNF-α (36, 37). The increased expression levels of TF and PAI-1 promoted VSMC proliferation, migration and thrombosis and subsequent intimal hyperplasia and restenosis progression. Inhibition of TF expression or silencing of PAI-1 expression markedly reduced the VSMC density in the neointima (38, 39). The present study showed that HGK decreased TF and PAI-1 expression in TNF-α-treated VSMCs by regulating the AKT pathway. In addition, similar phenomena were observed in denuded femoral arteries. Based on these data, HGK decreased VSMC proliferation, migration, and inflammation in vitro and neointimal hyperplasia in vivo, possibly partially by reducing TF and PAI-1 expression.

## 5. Conclusion

HGK inhibited the proliferation and migration of PDGF-BB-treated VSMCs and inflammation in TNF-α-stimulated VSMCs in vitro and ameliorated neointimal hyperplasia in the denuded femoral arteries in vivo. The antiproliferative, anti-migratory and anti-inflammatory effects of HGK on PDGF-BB- or TNF-α-stimulated VSMCs were mediated mainly by PDK1/AKT/mTOR/S6K. In addition, HGK promoted EPC chemotaxis and reendothelialization after injury. Thus, HGK may be a candidate treatment for intimal hyperplasia and restenosis.

## Abbreviations

VSMC: vascular smooth muscle cell
TNF-α: tumor necrosis factor-α
PDGF-BB: platelet-derived growth factor-BB
HGK: hydroxygenkwanin
EPCs: endothelial progenitor cells
VCAM-1: vascular cell adhesion protein 1
ICAM-1: intercellular adhesion protein 1
ATP7A: copper-transporting ATPase
Rac1: Rac family small GTPase 1
PDK1: phosphoinositide-dependent protein kinase 1
ED: endothelial denudation
PAI-1: plasminogen activator inhibitor 1
TF: tissue factor
PTEN: phosphatase and tensin homolog
mTOR: mammalian target of rapamycin
S6K: S6 kinase
PCNA: proliferating cell nuclear antigen
ROS: reactive oxygen species

## Ethics approval

This study was performed in line with the principles of the Declaration of Helsinki. This protocol was approved by the National Taiwan University College of Medicine and the College of Public Health’s Institutional Animal Care and Use Committee (IACUC 20150293).

## Acknowledgement

The graphic abstract was created in BioRender.com.

## Consent for publication

Not applicable.

## Data Availability

The authors confirm that the data supporting the findings of this study are available within the article and its supplementary materials.

## Author contributions

M.S.L., C.C.C. and Y.L.L.: Conceptualization, Methodology. Resources., P.Y.C.: Investigation, Visualization., S.H.W.: Investigation, Data curation, Writing-Original draft preparation. All authors reviewed, revised, and approved in the final manuscript.

## Supporting Information

**Supplemental table 1.** Antibodies used in the experiments.

| Antibody | Source | Code |
| --- | --- | --- |
| GAPDH | GeneTex | GTX100118 |
| β-actin | GeneTex | GTX112141 |
| α-tubulin | GeneTex | GTX109636 |
| phospho- PDK1 | GeneTex | GTX55111 |
| total PDK1 | GeneTex | GTX59809 |
| phospho-AKT (S473) | GeneTex | GTX128414 |
| phospho-AKT (T308) | GeneTex | GTX52321 |
| total AKT | GeneTex | GTX121937 |
| α-SMA | GeneTex | GTX100034 |
| BrdU | GeneTex | GTX28039 |
| PAI-1 | GeneTex | GTX100550 |
| P27 | GeneTex | GTX100446 |
| total mTOR | GeneTex | GTX101557 |
| total S6K | GeneTex | GTX107562 |
| Rabbit IgG isotype control antibody | GeneTex | GTX35035 |
| Mouse IgG isotype control antibody | GeneTex | GTX35009 |
| Cyclin E | Cell Signaling | #4129 |
| Cyclin D1 | Cell Signaling | #2978 |
| phospho- mTOR | Cell Signaling | #5536 |
| phospho- S6K | Cell Signaling | #9234 |
| CDK2 | Santa Cruz Biotechnology | sc-6248 |
| CDK4 | Santa Cruz Biotechnology | sc-56277 |
| VCAM-1 | Santa Cruz Biotechnology | sc-13506 |
| ICAM-1 | Santa Cruz Biotechnology | sc-8439 |
| E-selectin | Santa Cruz Biotechnology | sc-137054 |
| PCNA | Santa Cruz Biotechnology | sc-56 |
| ATP7A | LSBio | LS-C413152 |
| MMP-9 | Abcam | ab228402 |
| Cx43 | Abcam | ab230537 |
| Rac1 | Abcam | ab282581 |
| CD31 | Abcam | ab28364 |
| TF | BIOSS | bs-4690R |
| Na<sup>+</sup> K<sup>+</sup> -ATPase | arigo | ARG66327 |
| CD68 | arigo | ARG10514 |

**Supplemental table 2.**
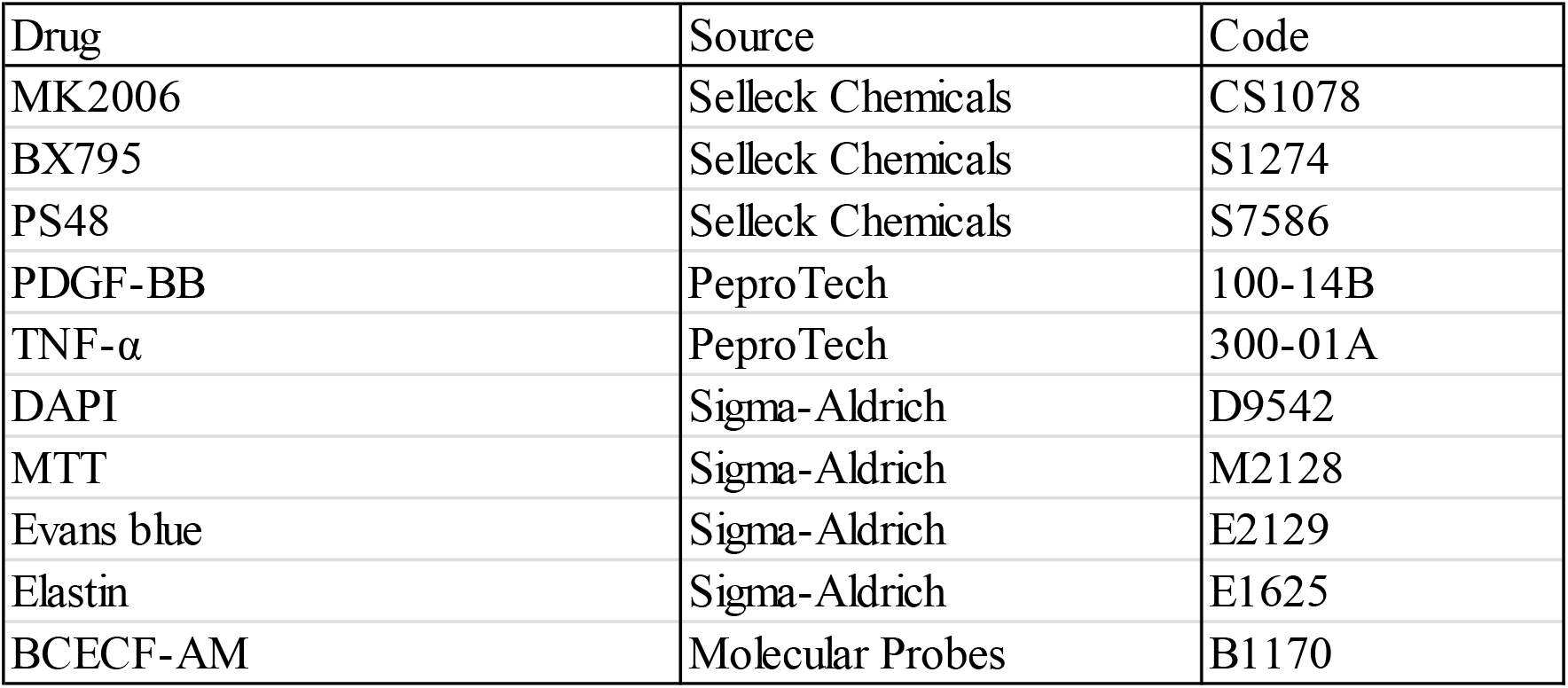
Drugs used in the experiments.

**Supplemental Figure 1.**
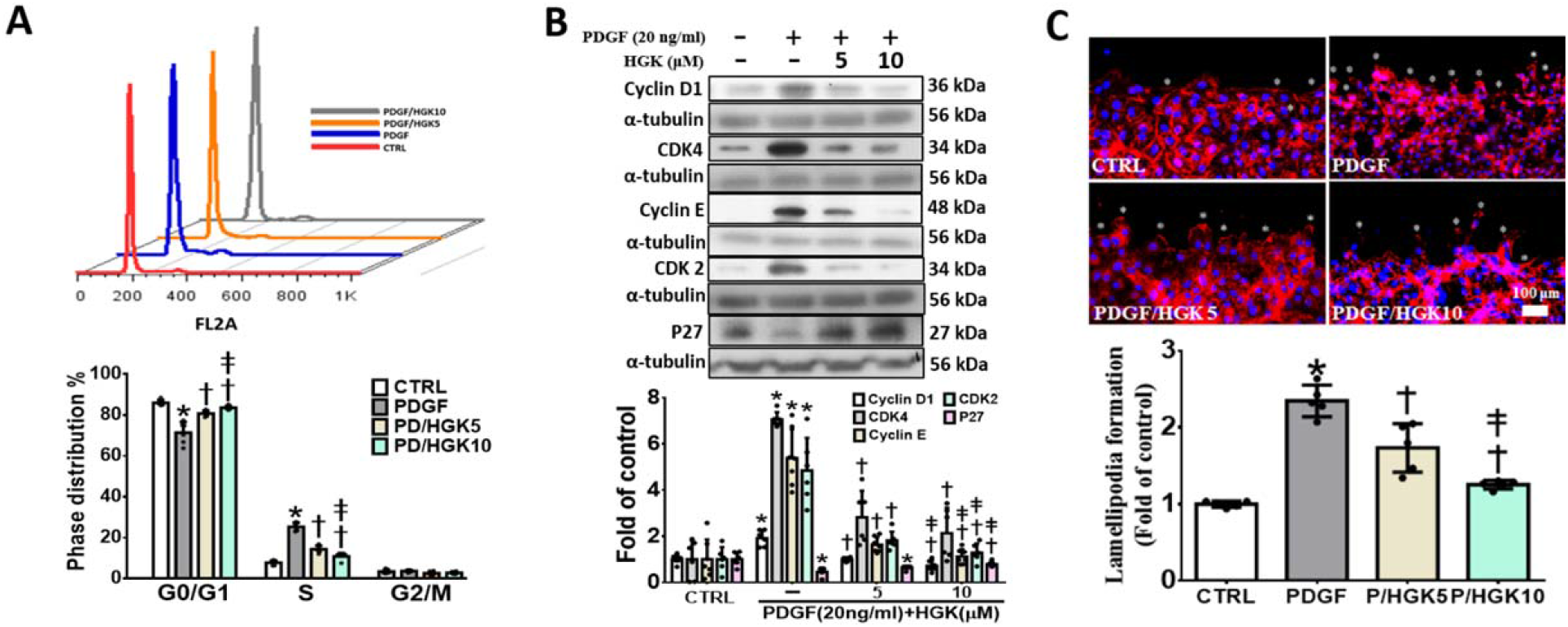
HGK decreases PDGF-BB-induced VSMC proliferation and migration. Serum-starved VSMCs were pretreated with 5 or 10 μM HGK for 1 h and were then treated with 20 ng/ml PDGF-BB (PDGF) for different times. (A-B) Flow cytometry and western blotting analysis showed that HGK decreased PDGF-induced VSMC proliferation and the expression levels of related cell cycle regulatory proteins. (C) Staining of actin fibers (phalloidin) and nuclei (DAPI) and their merged images are shown. Rhodamine-Phalloidin staining indicated that HGK decreased PDGF-induced VSMC lamellipodia formation at the leading edge. The scale bar represents 100 μm. Nuclei were stained with DAPI._The values are presented as the means ± SDs. *p < 0.05 compared with the CTRL group. †p < 0.05 compared with the PDGF group. ‡p < 0.05 compared with the PDGF/HGK5 group. N=5-12. CTRL, control group. Statistical comparisons were performed using one-way ANOVA.

**Supplemental Figure 2.**
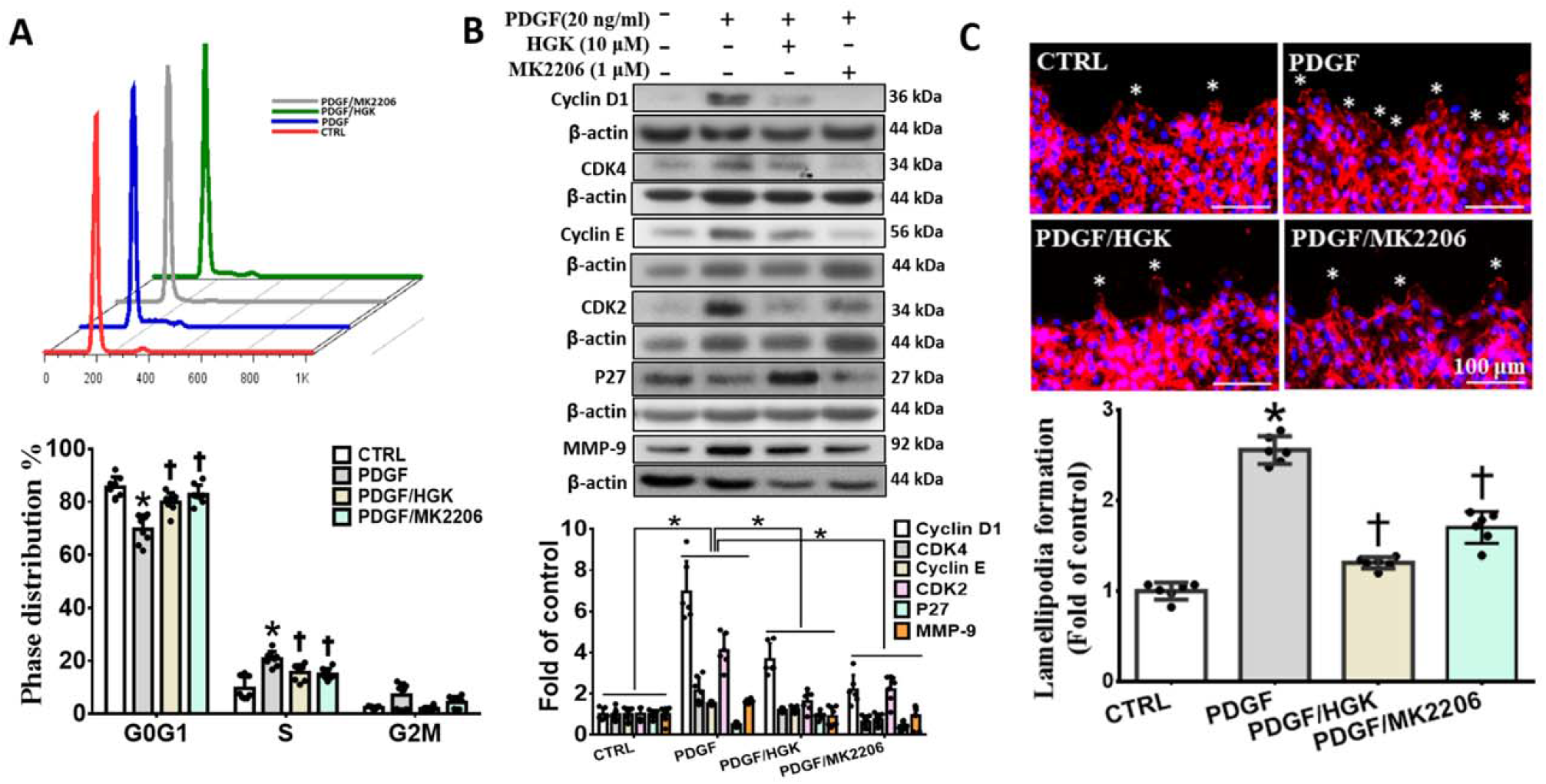
HGK decreases PDGF-BB-induced VSMC proliferation and migration by inhibiting the AKT pathway. Serum-starved VSMCs were pretreated with 10 μM HGK or 1 μM MK2206 (an AKT inhibitor) for 1 h and were then treated with 20 ng/ml PDGF-BB (PDGF) for different times. (A-B) Flow cytometry and western blotting analysis were performed, and these results showed that HGK decreased PDGF-induced VSMC proliferation and the expression levels of related cell cycle regulatory proteins by inhibiting AKT phosphorylation. (C) Rhodamine-Phalloidin staining indicated that HGK decreased PDGF-induced VSMC lamellipodia formation by inhibiting AKT phosphorylation. The scale bar represents 100 μm. Nuclei were stained with DAPI._The values are presented as the means ± SDs. *p < 0.05 compared with the CTRL group. †p < 0.05 compared with the PDGF group. ‡p < 0.05 compared with the PDGF/HGK5 group. N=5-12. CTRL, control group. Statistical comparisons were performed using one-way ANOVA.

